# The essential domain of FtsN triggers cell division by promoting interaction between FtsL and FtsI

**DOI:** 10.1101/2023.05.12.540521

**Authors:** Kyung-Tae Park, David Johnson Park, Sebastien Pichoff, Shishen Du, Joe Lutkenhaus

**Author notes:** To whom correspondence should be addressed: Joe Lutkenhaus, Department of Microbiology, Molecular Genetics and Immunology, University of Kansas Medical Center, Kansas City, Kansas, USA.

## Abstract

Cell division in bacteria requires the activation of FtsWI at the division site to synthesize septal peptidoglycan. In *E. coli* FtsN activates FtsWI and a previous model posited that the essential domain of FtsN (^E^FtsN) acts on FtsQLB causing conformational changes so that a domain of FtsL, called AWI (^AWI^FtsL), contacts FtsI resulting in activation of FtsW. In this study we use genetic analysis along with an AlphaFold2 model to test this activation model. Based on our findings we propose an updated model wherein the ^AWI^FtsL and FtsI interaction is stabilized by ^E^FtsN to activate FtsW and that this interaction is enhanced by the ^Cyto^FtsN-FtsA interaction. Thus, FtsN acts as both a sensor for divisome assembly and an activator. In addition, we elucidate the role played by two critical FtsL residues in FtsW activation.

## Introduction

A major step in bacterial cell division is the activation of septal peptidoglycan (sPG) synthesis at the Z ring, a cytoskeletal element that is formed by the coalescence of membrane-attached FtsZ filaments at midcell ^1–3^. In *E. coli*, FtsZ filaments attach to the membrane by interacting with the membrane tethers FtsA and ZipA ^4, 5^. The Z ring serves as a scaffold with FtsA serving as a hub for recruitment of the other division proteins including the glycosyltransferase (GT) FtsW and its cognate transpeptidase (TP) FtsI, which together synthesize sPG leading to constriction of the cell envelope ^6, 7^. These proteins are recruited by the FtsQLB complex and once assembled, FtsN arrives and activates the FtsWI enzyme complex by turning FtsQLB from a recruitment complex to an activation complex ^8, 9^.

Genetic evidence suggests FtsN provides an activation signal on both sides of the membrane ^9^. FtsN is a single pass transmembrane protein and contains two important domains that are required for it to function as an activator. A short cytoplasmic domain (^Cyto^FtsN) interacts with FtsA while a periplasmic essential domain (^E^FtsN), consisting of a ∼dozen amino acids (80-92) embedded in a lengthy disordered region, acts on the FtsQLB complex ^9,10^. At physiological levels both interactions are required, however, ^E^FtsN is sufficient to activate constriction when overproduced and exported to the periplasm ^11^. In addition to ^Cyto^FtsN and ^E^FtsN, the carboxy terminus of FtsN contains a SPOR domain (^SPOR^FtsN) that binds to denuded glycan chains produced during septation ^11, 12^. This interaction is not essential but enhances the efficiency of FtsN localization by generating a positive feedback loop and is required for proper septal morphology ^11, 13^.

FtsN requires all other essential division proteins to localize to the division site and is thought to be the trigger for initiation of sPG synthesis ^11, 14, 15^. This multiprotein requirement for the localization of FtsN is due in part to ^Cyto^FtsN binding to FtsA and the ^E^FtsN interacting with FtsQLB ^9, 10^. However, FtsW and FtsI are also required ^15^. Support for FtsN being the trigger for septal PG synthesis also comes from tracking molecules *in vivo*. FtsWI complexes moves at two different speeds at midcell ^7^. An inactive complex moves at a faster speed that is dictated by the treadmilling of FtsZ filaments. In contrast, an active complex moves at a slower speed, is disengaged from FtsZ filaments and is actively synthesizing peptidoglycan. FtsN is only seen moving at the slower speed suggesting that the arrival of FtsN at midcell switches FtsWI from an inactive to an active complex ^16^. This is consistent with the evidence that FtsN is the trigger for the initiation of constriction.

A model for activation of FtsWI was proposed based on mutations that either activate (superfission mutations) or block septal PG synthesis ^8^. Dominant negative mutations (which are also loss of function mutations [LOF]) in *ftsL* defined a region of FtsL, designated AWI (<u>A</u>ctivation of Fts<u>W</u> and FtsI), required for activation of FtsWI. Of note, no dominant negative mutations were isolated in *ftsB* or *ftsQ*, underscoring the critical and unique role of FtsL in activation despite it forming a tight complex with FtsB and FtsQ ^17^. The strongest of these mutations *are ftsL^L86F^* and *ftsL^E87K^*. Since these mutations are resistant to the action of FtsN, it was proposed that ^E^FtsN causes a conformational change in FtsQLB that allows ^AWI^FtsL to interact with FtsI, which activates FtsW. In contrast, superfission mutations in *ftsL* and *ftsB* require less (or bypass) FtsN and define regions in FtsL and FtsB designated CCD (<u>C</u>onstriction <u>C</u>ontrol <u>D</u>omain) ^9^. Since the strongest of these mutations (*ftsB^E56A^* and *ftsL^E88K^*) have lost or swapped charge, it is likely that they disrupt an interaction suggesting that ^E^FtsN alters the CCD domains and that the CCD superfission mutations mimic ^E^FtsN action. *In vitro* reconstitution using proteins isolated from *Pseudomonas aeruginosa* revealed that FtsQLB or just FtsLB stimulates the GT activity of FtsW ^18^. Consistent with the model, activation was blocked by some of the dominant negative mutants of FtsL. Since the activation *in vitro* was not stimulated by FtsN, it suggested that the isolated FtsQLB complex was in the activated state but at least some of the dominant negative *ftsL* mutations locked the complex in the inactive state.

Recently, AlphaFold2 (AF2) was used to predict the structure of the FtsQLBWI complex which suggested ^AWI^FtsL is in contact with FtsI ^19^. In a another study FtsN was modeled along with this complex and the only region of FtsN making significant contact with the FtsQLBWI complex was ^E^FtsN, which contacts ^AWI^FtsL and FtsI ^20^. Previously, we suggested that a mutation altering ^AWI^FtsL (*ftsL^E87K^*) would interfere with FtsL interacting with FtsI ^8^. In part, this was based on the failure of a strong *ftsB* superfission mutation (*ftsB^E56A^*), which was reported to bypass FtsN ^9^, to suppress *ftsL^E87K^*. However, we subsequently found that the bypass of *ftsN* by *ftsB^E56A^*was media dependent and did not occur in rich media which was used in our previous study. Since the AF2 model suggests that ^AWI^FtsL is part of the binding site for ^E^FtsN, we investigated the significance of ^E^FtsN’s modeled interaction with ^AWI^FtsL and FtsI and the roles of key FtsL residues in activation of FtsW. In our revised model (Fig. 1) the localization of FtsQLB and FtsWI to the Z ring leads to ^AWI^FtsL and FtsI forming a transient interface for ^E^FtsN binding that, along with ^Cyto^FtsN binding to FtsA, recruits FtsN to the divisome resulting in activation of FtsW with the activation signal going from ^E^FtsN through FtsI to FtsW.

**Fig. 1.**
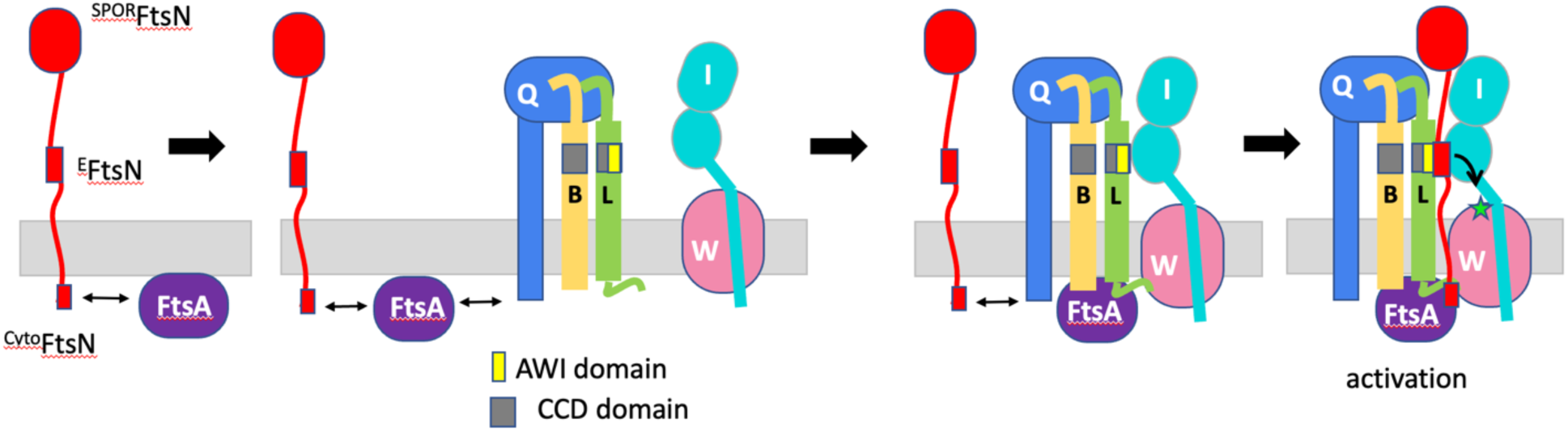
An updated model for role of ^E^FtsN in the activation of FtsWI based on this study. In this model ^Cyto^FtsN interacts with FtsA, but this interaction is transient. FtsA recruits the FtsQLB complex (with FtsK [not shown] as in intermediary) which recruits the FtsWI complex. Once the FtsQLBWI complex is assembled on FtsA, FtsN interacting with this FtsA through ^Cyto^FtsN facilitates ^E^FtsN binding to a site in the periplasm where ^AWI^FtsL weakly contacts FtsI. The binding of ^E^FtsN brings ^AWI^FtsL and FtsI closer together resulting in FtsL-L86 signalling to FtsI which is propagated to FtsW. Superfission mutations in *ftsL* and *ftsB* mimic the binding of ^E^FtsN resulting in a stronger interaction between ^AWI^FtsL and FtsI. This stronger interaction produces an activation signal which can still be enhanced by FtsN. Substitutions at L86 of FtsL prevent superfission mutations in *ftsL* and *ftsB* from propagating a signal to FtsI whereas the FtsL-L87K substitution prevents ^E^FtsN binding.

## Results

### The FtsQLBWIN complex

The predicted structure of the FtsQLBWI complex from *E. coli* obtained with AF2 is similar to that obtained by others ^19–21^(Fig. S1A). As noted, this model has high confidence and agrees with known interactions of the proteins in the complex. In *E. coli* the cytoplasmic domain of FtsL is required for recruitment of FtsW and in the predicted structure FtsL residues L24 and I28, required for recruitment of FtsW, are packed against hydrophobic residues in FtsW ^8^. Similar to the RodA/Pbp2 complex, the transmembrane domain of FtsI contacts two of the transmembrane helices of FtsW ^22^. Furthermore, FtsL and FtsB form a coiled coil that extends from the cytoplasm to the CCD regions of each protein while the C-terminal regions of FtsB, FtsL and FtsI form an extended beta-sheet with FtsQ, consistent with the structure of the FtsB-FtsQ and FtsQLB complexes ^23, 24^. Importantly, the residues that constitute ^AWI^FtsL are packed against FtsI.

In the complex, L86 of FtsL is docked in a hydrophobic pocket composed of residues V84, V86, Y168 and P170 from FtsI (Fig. 2A). Interestingly, the *ftsI^V86E^*mutation affects FtsN recruitment and FtsI function ^25^. Also, *ftsL^L86F^* is a strong dominant negative mutation and, as modeled, the phenylalanine residue would be too large to pack into this pocket. Consistent with this, we found that other drastic substitutions at this position (*ftsL^L86A^*, *ftsL^L86E^, ftsL^L86K^*) are also dominant negative mutations whereas a conservative substitution (*ftsL^L86V^*) behaves like WT indicating the importance of the size and hydrophobicity of the residue at this position (Fig. S2). Importantly, the *ftsL^L86F^* mutation did not affect recruitment of FtsI, presumably because FtsW is recruited before FtsI in the assembly pathway and the FtsI transmembrane domain, which is intimately associated with the TMs of FtsW (Fig. S1), is sufficient for FtsI recruitment ^8, 26^. This is consistent with *ftsL^L86F^*blocking a step after recruitment, likely preventing activation of FtsWI. Also, consistent with this, we previously found that superfission alleles of *ftsW*, *ftsW^M^*^269^*^I^* and *ftsW^E^*^289^*^G^* that are already activated, suppress *ftsL^L86F^* ^8, 27^.

**Fig. 2.**
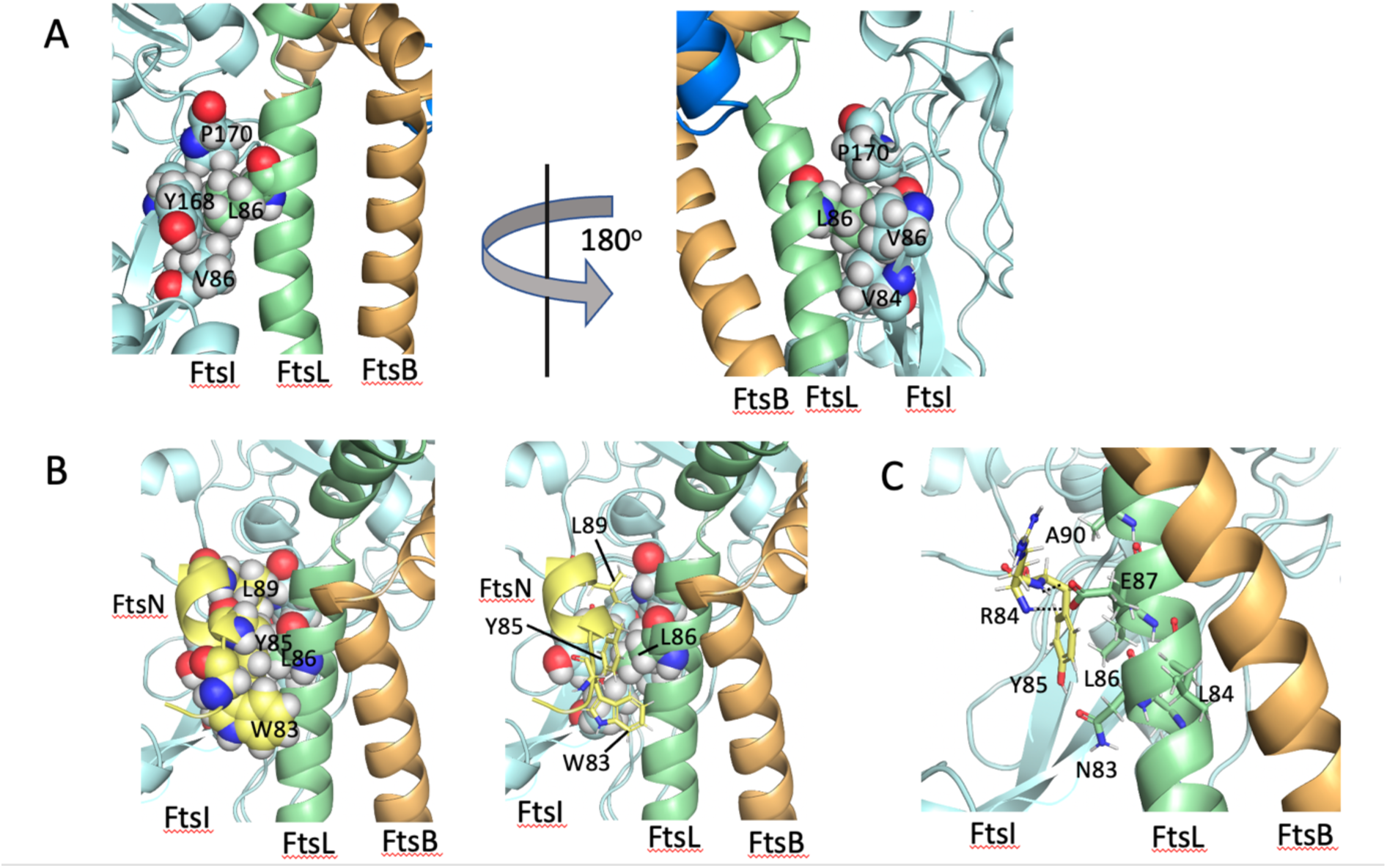
FtsL-L86 interacts with FtsI and ^E^FtsN whereas FtsL-E87 only interacts with ^E^FtsN. A) FtsL-L86 is embedded in a hydrophobic pocket formed by four FtsI residues. In the AlphaFold2 (AF2) model, residue L86 of FtsL is surrounded by hydrophobic residues from FtsI. The residues include P170, Y168, V86 and V84. B) The E domain of FtsN (^E^FtsN) binds to both FtsL and FtsI. In the AF2 model, ^E^FtsN binds to ^AWI^FtsL and FtsI, essentially sitting on top of L86 of FtsL. The three critical residues of ^E^FtsN are W83, Y85 and L89 with L89 being the most conserved residue. L89 and Y85 residues of ^E^FtsN in combination with the 4 hydrophobic residues from FtsI (Panel 2A) completely engulf FtsL-L86. In addition to contacting FtsL-L86 and A90, FtsN-L89 also contacts FtsI-P170 and Y168. In the right model the three residues of ^E^FtsN are depicted as sticks to better see how they interact with FtsL-FtsI. C) FtsL-E87 forms hydrogen bonds to backbone amides of residues in ^E^FtsN. Although E87 of FtsL does not interact with side chains of ^E^FtsN, it forms hydrogen bonds with the backbone amides of ^E^FtsN residues R84 and Y85. It does not contact FtsI.

When FtsN was modeled with this complex, a 69-residue segment of FtsN (75-143) was confidently predicted to interact primarily with FtsI, much of which had a pLDDT greater than 70, with only two regions having a defined secondary structure (Fig. S1B). The most functionally significant interaction involved ^E^FtsN, which was modeled as a short helix binding to FtsI and FtsL (Fig. 2B). This interaction is the only modeled region to interact with anything other than FtsI and involves the residues that constitute ^AWI^FtsL including L86 (Fig.S3). Mutations in this region of *ftsL* are resistant to activation by FtsN^8^.

Importantly, the modeled interaction between FtsN and FtsI and FtsL involves the three residues of FtsN determined to be critical for ^E^FtsN function, W83, Y85 and L89 ^9^(Fig. 2B). Analysis of FtsN from a variety of Gram-negative bacteria indicate that the consensus sequence for ^E^FtsN is (WYLF)X(YFL)hXXL with L89 being completely conserved (Fig. S4). Y85 and L89, along with the aforementioned four hydrophobic residues from FtsI, complete the engulfment of the FtsL-L86 residue further constraining the type of residue that could fit in this pocket. In addition to contacting FtsL-L86, FtsN-Y85 contacts FtsI-Y168 and FtsN-L89 contacts FtsI-P170 (Fig. 2B). The other key ^E^FtsN residue W83 packs against hydrophobic residues in FtsL, L84 and N83, both of which are important for FtsL function and result in a dominant negative phenotype when mutated ^8^(Fig. 2B and S3). Two additional FtsL residues produce a dominant negative phenotype when mutated, A90 and E87 ^8^. FtsL-A90 abuts FtsN-L89 and the *ftsL^A90E^* mutation would interfere with FtsN binding due to the introduction of the larger glutamate residue (Fig. 2C). The side chain of FtsL-E87 is extended to form two hydrogen bonds with the backbone amides of R84 and Y85 of FtsN, which together helps form the binding pocket for residues W83 and Y85 of FtsN (Fig. 2C). Previous mutagenesis has shown that loss of the negative charge at FtsL-E87 results in dominant negative phenotype ^8^. The AF2 model suggests that it is the loss of two hydrogen bond acceptors present on a negatively charged residue, rather than loss of the negative charge itself, that affects the binding of ^E^FtsN.

Superfission mutations in *ftsL* and *ftsB* are thought to mimic FtsN action ^8, 9, 28^. Two of the stronger mutations are *ftsL^E88K^* and *ftsB^E56A^*. In the AF2 model these two residues interact with FtsB^R70^ forming a network of charged groups (Fig. S5). Since removal of the negative charge by mutation (*ftsL^E88K^* and *ftsB^E56A^*) results in a superfission phenotype it suggests that disruption of this charge network results in conformational changes leading to activation. Consistent with this, the *ftsB^R70C^* mutation, which would also disrupt this charge network, is also an activation mutation ^29^. *ftsL^G92D^* is also a superfission mutation and the introduction of the aspartic acid side chain into this region would also disrupt the charge network as it would cause a steric clash with R70.

In the AF2 model, a region of FtsN located around FtsN^117–123^ is modeled as a helix that interacts with FtsI (Fig. S7). This is the location of a putative helix 3 within the disordered region of FtsN that extends from the membrane to the SPOR domain ^11^. To test the importance of this interaction, we made an FtsN mutant containing three mutations in the most conserved residues in this region. This triple mutant was not affected in a complementation assay (Fig. S6B). However, adding these three mutations to *ftsN^D5N^*, resulted in two-fold more IPTG being required for complementation compared to just *ftsN^D5N^*. The *ftsN^D5N^*mutation reduces the interaction of FtsN with FtsA so that FtsN has to be increased several fold for it to complement a 11*ftsN* strain ^10^. Thus, adding the three mutations to a less active mutant (*ftsN^D5N^*) displays a slight phenotype indicating this region may make a minor contribution to FtsN function.

### FtsL^E87K^ is refractory to ^E^FtsN

The AF2 model indicates that ^E^FtsN not only binds to FtsL and FtsI, but also brings them closer together. Using FtsW as a reference, the whole TP domain of FtsI is angled closer to FtsL, while FtsL, FtsB, and FtsW are all pushed a little away, probably to accommodate FtsN (Fig. S7). Furthermore, the model indicates that the mutations in *^AWI^ftsL* should abrogate ^E^FtsN binding. For the *ftsL^E87K^* mutation this may be the only effect as E87 only interacts with the backbone amides of FtsN and does not contact FtsI (Fig. 2B). In contrast, the *ftsL^L86F^* mutation would likely affect binding to FtsI as well as FtsN as residues L89 and Y85 of FtsN make hydrophobic contacts with L86 of FtsL (Fig. 2B). This suggests that these two mutations, *ftsL^E87K^* and *ftsL^L86F^*, may affect different steps in FtsWI activation and we set out to test this by examining the effect of these mutations on the activation of FtsW by ^E^FtsN.

To test the effect of *ftsL^E87K^* on the activation of FtsWI by ^E^FtsN we utilized a strain that grew in the absence of FtsN but still responded to ^E^FtsN. We used the *ftsW^E^*^289^*^G^* allele as it bypasses FtsN more efficiently than *ftsW^M^*^269^*^I^*, *ftsL^E88K^*, *ftsB^E56A^* and *ftsI^K^*^211^*^I^* mutations as cells are only mildly longer, even in rich media, in the absence of FtsN ^9, 27, 28^. We confirmed that *ftsB^E56A^* also bypasses FtsN, but we only obtained this bypass on minimal media and cells are more filamentous on rich media (consistent with our previous study showing *ftsB^E56A^* is unable to rescue *ftsL^E87K^* on rich media ^8^). Although *ftsW^E^*^289^*^G^* bypasses FtsN, it is not fully active as it still responds to FtsN resulting in a smaller cell size. We set up a test to determine if *ftsL^E87K^* negates responsiveness to FtsN.

For the test we used a strain containing *ftsW^E^*^289^*^G^* on the chromosome and deleted *ftsN* by transduction of an *ftsN::kan* allele. After removal of the Kan resistance marker, we introduced plasmids carrying WT *ftsL*, *ftsL^E87K^*, *ftsL^E88K,G92D^*, or *ftsL^E87K,E88K,G92D^*under arabinose promoter control. We included *ftsL^E88K,G92D^* as we showed previously that these mutations are synergistic; the combination of these two mutations rescues FtsL truncated for its cytoplasmic domain whereas the single mutations do not ^8^. FtsL truncated for its cytoplasmic domain is unable to recruit FtsWI, however, recruitment is restored by the combination of these superfission mutations. In full length *ftsL* these two mutations weakly bypass FtsN indicating they act downstream of ^E^FtsN action (Fig. S8). We next transduced in *ftsL::kan* in the presence of arabinose. Transductants were obtained regardless of the *ftsL* allele on the plasmid which was expected since we previously showed that *ftsW^E^*^289^*^G^* suppresses *ftsL^E87K^*^8, 27^. Consistent with expectations, fewer transductants were obtained when the strain carried *ftsL^E87K^*compared to WT *ftsL*. The reduced frequency in the presence of *ftsL^E87K^* was expected since *ftsW^E^*^289^*^G^* on the chromosome would be replaced by WT *ftsW* by contransduction (about 70%). Such cotransductants cannot grow in the presence of *ftsL^E87^*^K^. Interestingly, the number of transductants was not reduced in the presence of *ftsL^E88K,G92D^*or *ftsL^E87K,E88K,G92D^* indicating that *ftsL^E87K^*is not deleterious in the presence of the two superfission mutations. To be sure of the strains, we checked by PCR to ensure that *ftsL::kan* had replaced *ftsL* on the chromosome and the plasmids retained the *ftsL* mutations (Fig. S8A). Interestingly, the strains carrying *ftsL^E88K,G92D^* or *ftsL^E87K,E88K,G92D^* produced a smaller cell phenotype than the strains carrying *ftsL* or *ftsL^E87K^*alleles (Fig. 3A). This suggests that the two activation mutations enhance the activity of the *ftsW^E^*^289^*^G^* strain even in the absence of *ftsN* and that *ftsL^E87K^* does not block this activity. This is consistent with E87 of FtsL only interacting with FtsN.

**Fig. 3.**
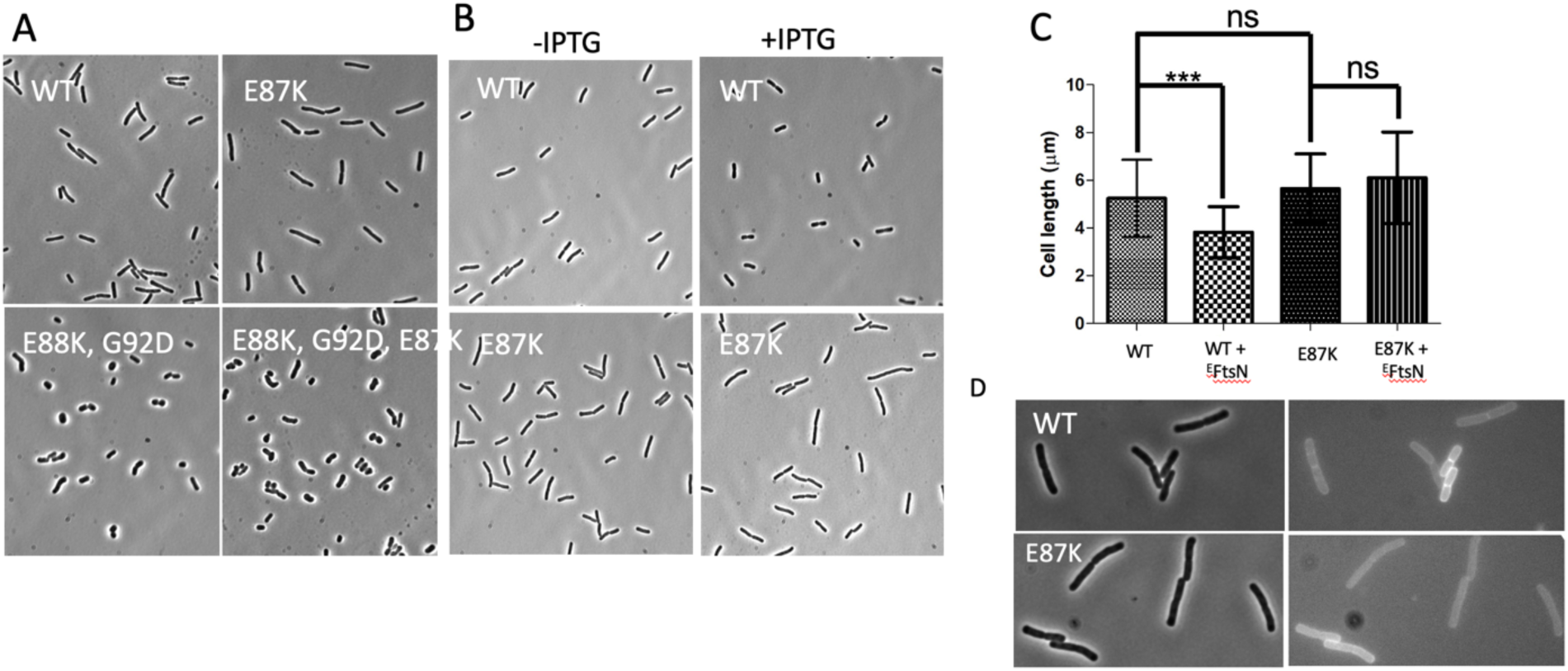
FtsL^E87K^ is nonresponsive to ^E^FtsN. A) PK4980-1 (*ftsW^E289G^*, Δ*ftsN*, *ftsL::kan*) containing plasmids expressing *ftsL*^WT^, *ftsL^E87K^*, *ftsL^E88K,G92D^* or *ftsL^E88K,G92D,E87K^* under arabinose promoter control (derivatives of pSD296 [P_ara_::*ftsL*]) were grown to exponential phase in liquid medium and photographed. B) The strains in A containing plasmids expressing *ftsL* or *ftsL^E87K^* were transformed with pMG14 *(bla^R^,* P_lac_::*^TT^gfp-ftsN*^71^–*^1^*^05^ [E domain]) or an empty vector. Purified transformants were grown overnight and diluted 1:200 into the same medium and grown at 37°C with the addition of 200 µM IPTG to induce GFP-^E^FtsN. When the OD_600_ reached 0.3, samples were taken for microscopy. C) The average cell lengths of the cells from the cultures photographed in B were determined (N>200 per each culture) as described in Materials and Methods. D) Strains as in B but induced with 50 μM IPTG. Cells were photographed 1 hr after induction at 30°C with phase on the left and fluorescence on the right.

To further test the responsiveness of FtsL^E87K^ to ^E^FtsN, we introduced a plasmid which has ^E^FtsN with a Tat export signal fused to GFP under control of the *lac* promoter (pMG14, ^11^) into strains expressing WT *ftsL* or *ftsL^E87K^*. The strains were grown to exponential phase with or without IPTG and the cell lengths determined to see if expression of ^E^FtsN reduced the average cell length. Although *ftsW^E^*^289^*^G^* bypasses *ftsN*, strains carrying this mutation still respond to FtsN by a reduction in average cell length as observed above with the superfission mutations in *ftsL*. In the absence of ITPG the strains with *ftsL* or *ftsL^E87K^* had a similar average cell length (*ftsL^WT^* 5.2+/-1.6 vs *ftsL^E87K^* 5.6+/-1.45) (Fig. 3B & C). In contrast, in the presence of IPTG the strain containing WT FtsL displayed a significant reduction in the average cell length (27% decrease) indicating it responded to ^E^FtsN. In contrast, IPTG did not significantly affect the average cell length of the strain with *ftsL^E87K^*. Thus, we conclude *ftsL^E87K^* is resistant to ^E^FtsN.

Since the strain expressing *ftsL^E87K^* failed to respond to ^E^FtsN-GFP, a simple possibility is that this mutation prevents interaction with ^E^FtsN-GFP as suggested by the AF2 model (Fig. 2C). Although ^E^FtsN-GFP localization cannot be detected in WT cells ^11^, it can be detected in cells carrying a superfission mutation (*ftsB^E56A^*) suggesting such mutations may result in a higher affinity of ^E^FtsN for the divisome. Since our strains carry a superfission mutation (ftsW^E289G^), it may allow ^E^FtsN-GFP localization to be detected. To test this, strains carrying *ftsL* or *ftsL^E87K^* were grown in liquid media and examined for ^E^FtsN-GFP localization one hour after induction with IPTG. Indeed, we could readily detect localization of ^E^FtsN-GFP to the septum in cells expressing *ftsL* (Fig. 3D). In contrast, we failed to detect localization in cells expressing *ftsL^E87K^* (Fig. 3). We conclude that *ftsL^E87K^* prevents the binding of ^E^FtsN.

### FtsL L86 is required for activation of FtsW

In the current model for activation of FtsWI, ^E^FtsN acts on FtsQLB complex resulting in a conformational change in FtsQLB allowing ^AWI^FtsL to interact with FtsI which then signals to FtsW ^8^. Based on the AF2 model, both residues L86 and E87 of FtsL are involved in binding to ^E^FtsN, however, FtsL-L86 is also involved in binding FtsI whereas FtsL-E87 only makes contact with ^E^FtsN. This suggests that mutations altering FtsL-L86 would be more disruptive than mutations affecting FtsL-E87 as they would also affect interaction with FtsI and could prevent the transmission of the activation signal to FtsI. One way to test this is to examine the effects of superfission mutations in *ftsL* and *ftsB* that are thought to mimic the effects of the action of ^E^FtsN. If the *ftsL* mutations only disrupt the interaction with FtsN, the superfission mutations in *ftsL* or *ftsB* may rescue the dominant negative mutations in *ftsL* (since these superfission mutations require less FtsN). On the other hand, if the *ftsL* mutations affect interaction with FtsI, and this is needed for activation of FtsW, then the superfission mutations would not help (or have any effect) as the activation signal would be blocked and not delivered through FtsI to FtsW. If so, such mutations should be rescued by *ftsW^E^*^289^*^G^* as it suppresses both *ftsL^E87K^*and *ftsL^L86F^* and the activation signal from FtsN is not required ^8^.

As a test of the model, we checked whether superfission mutations in *ftsL* could suppress the dominant negative *ftsL* mutations. To do this, the superfission mutations in *ftsL* were introduced in *cis* with *ftsL^E87K^* and tested for complementation of an FtsL depleted strain (Fig. 4A). Although *ftsL^E88K^* could not suppress *ftsL^E87K^*, *ftsL^G92D^* did suppress *ftsL^E87K^*as growth was restored. *ftsL^G92D^* also suppressed *ftsL^A90E^* as expected as it is a weaker dominant negative mutation than *ftsL^E87K^*and it is also rescued by *ftsL^E88K^* ^8^. Since *ftsL^G92D^* suppressed *ftsL^E87K^* we checked to see if it could suppress *ftsL^L86F^*, however, it did not suppress *ftsL^L86F^* (Fig. 4A). It could not suppress *ftsL^L86E^*or *ftsL^L86K^* either. This result argues that *ftsL^G92D^* can compensate for the loss of FtsL-E87’s ability to bind ^E^FtsN but cannot compensate for the effect *fts^L86F^*has on the complex.

**Fig. 4.**
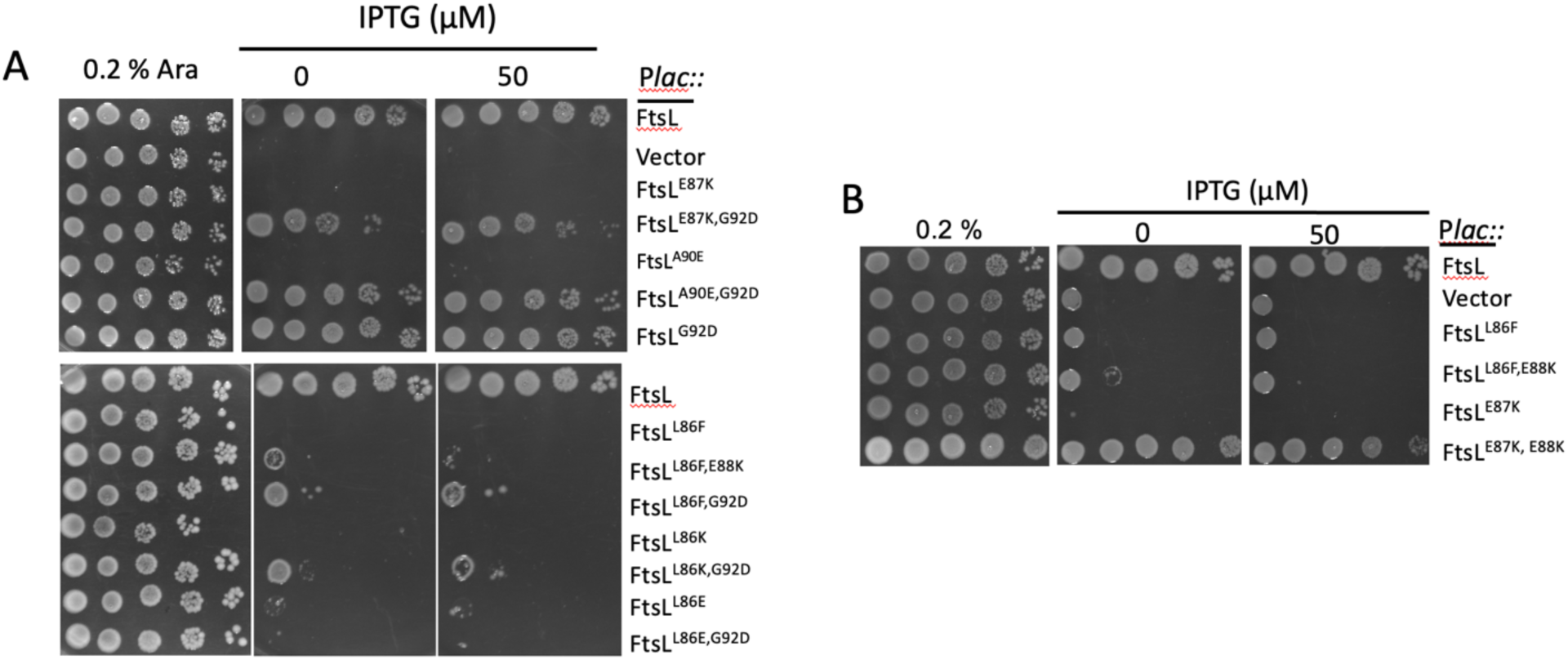
Superfission mutations in *ftsL* and *ftsB* suppress *ftsL^L87K^* but not *ftsL^L86F^*. A) *ftsL^G92D^* suppresses *ftsL^L87K^* but not *ftsL^L86F^*. PK439-2 (*ftsL::kan*/pSD296 [P_ara_::*ftsL*]) *recA1*) was transformed with pKTP100 (*bla^R^*, P_tac_::*ftsL*) or derivatives containing various alleles of *ftsL*. The strains were spotted on plates containing 0.2% arabinose (positive control) or plates lacking arabinose and containing various concentrations of IPTG and incubated at 37°C. B) Combining *ftsB^E56A^ in trans* and *ftsL^E88K^* in *cis* suppresses *ftsL^E87K^* but not *ftsL^L86F^*. PK169-1 (*ftsB^E56A^*, *recA::spc, ftsL::kan*/ pDS296 [P_ara_::*ftsL*]) was transformed with pKTP100 (*bla^R^*, P_tac_::*ftsL*) carrying various *ftsL* alleles and grown in 0.2% arabinose and the necessary antibiotics. The strains were spotted on plates containing 0.2% arabinose or on plates lacking arabinose and containing various concentrations of IPTG and incubated at 37°C.

As shown above *ftsL^G92D^* suppressed *ftsL^E87K^*, but *ftsL^E88K^* did not. In addition, *ftsB^E56A^* did not suppress *ftsL^E87K^* (consistent with *ftsB^E56A^* being unable to bypass FtsN in rich media). To see if the ^E^FtsN-mediated signal goes through FtsL to FtsI (and to FtsW) and requires the interaction of ^AWI^FtsL with FtsI, we tested synergism between these superfission mutations in suppressing *ftsL^E87K^*. The combination of *ftsL^E88K^*and *ftsB^E56A^* suppressed *ftsL^E87K^*, however, neither *ftsL^E88K^* nor *ftsB^E56A^* or the combination of these two superfission mutations suppressed *ftsL^L86F^*(Fig. 4B). Taken together, these results indicate that the activation signal produced by *ftsL^G92D^*or *ftsL^E88K^* in combination with *ftsB^E56A^* is able to compensate for the loss of ^E^FtsN interaction to activate FtsWI. However, these superfission mutations cannot suppress *ftsL^L86F^* indicating the activation signal is not transmitted to FtsI. This supports a model where the activation signal goes from ^E^FtsN to ^AWI^FtsL involving FtsL-L86 that interacts with FtsI to activate FtsW.

### FtsI-Y168 involvement in activation of FtsW

The above results show that FtsL-L86 is required to interact with FtsI to activate FtsW. This was true whether the activation signal was due to ^E^FtsN or was due to superfission mutations in *ftsL* and *ftsB* that mimic ^E^FtsN action. In either case the *ftsL^L86F^*mutation prevented activation of FtsW. However, when an activated version of FtsW was present (FtsW^E289G^) *ftsL^L86F^* was rescued as the activation signal was no longer required. Since FtsL-L86 interacts with several residues in FtsI a prediction is that changing these residues would block activation by ^E^FtsN but would be rescued by *ftsW^E^*^289^*^G^*provided the substitutions in FtsI did not interfere with the TP activity of FtsI. To test this idea, we made substitutions at FtsI-Y168, one of four hydrophobic residues that contacts FtsL-L86 and tested them for complementation and for rescue by *ftsW^E289G^*.

Two mutations were made in *ftsI*, *ftsI^Y168A^* and *ftsI^Y168K^*. These alleles were introduced into a FtsI temperature-sensitive (Ts) strain and tested for their ability to complement the Ts mutation. Although *ftsI^Y186A^* partially complemented this strain (cells were filamentous), *ftsI^Y168K^* did not suggesting that the FtsI-Y168 residue is important in activating FtsW by ^E^FtsN acting through ^AWI^FtsL (Fig. 5). To test for rescue of *ftsI^Y168K^* by *ftsW^E289G^*, which would also rule out instability of FtsI^Y186K^, a plasmid expressing *ftsW^E289G^*under *lac* promoter control was introduced into these strains. This plasmid expresses sufficient FtsW^E289G^ to complement a Δ*ftsW* strain in the absence of added IPTG ^30^. The presence of *ftsW^E289G^*rescued *ftsI^Y186K^* indicating that interaction between FtsI-Y168 and FtsL-L86 was no longer required and that the TP activity of FtsI^Y168K^ was not compromised (Fig. 5). Together, these results demonstrate that interaction between FtsL and FtsI in the region corresponding to FtsL-L86 and FtsI-Y168 is required for activation of FtsW but not for FtsI to function if FtsW is already activated.

**Fig. 5.**
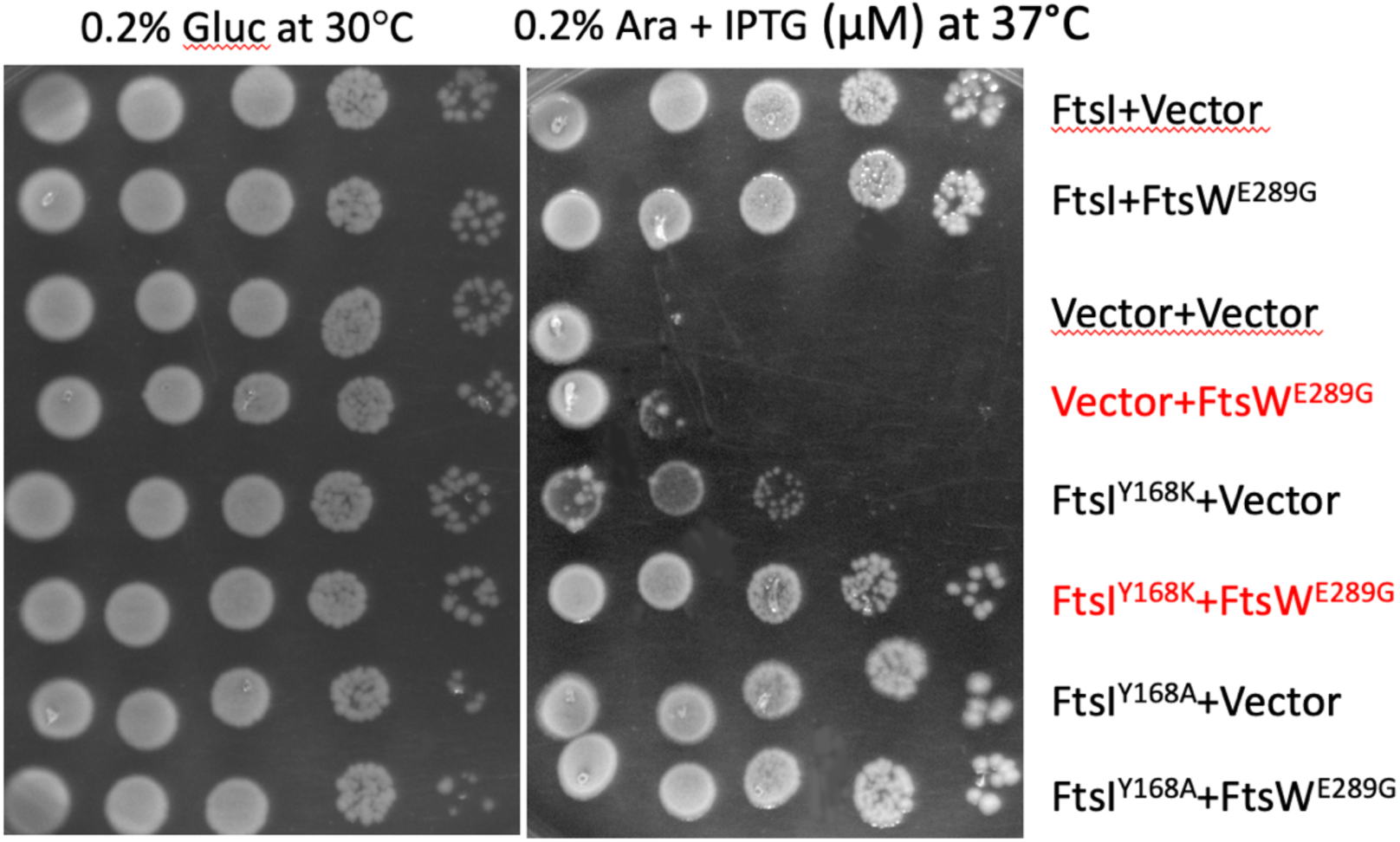
FtsI-Y168 is required for propagation of the activation signal to FtsW. A) PK23 (*ftsI*^Ts]^ *recA::spc*) was transformed with pKTP109 (P*ara*::*ftsI*) or derivatives containing various *ftsI* mutations. The resultants strains were transformed with a second plasmid expressing *ftsW^E289G^* (pSEB420 [P*lac*:: *ftsW^E289G^*]) or a vector. The strains were spotted on plates with 0.2% arabinose (to induce the *ftsI* derivatives) and incubated at restrictive temperature (37°C) with a control plate at 30°C. Note that sufficient *ftsW^E289G^* is expressed in the absence of IPTG to complement *11ftsN*

## Discussion

In this study we used genetic analysis in combination with an AF2 model to investigate the role of ^E^FtsN in the activation of FtsWI during cell division. The results lead to a revised model (Fig. 1) in which the recruitment of FtsQLB and then FtsWI to an FtsA leads to a weak interaction between ^AWI^FtsL and FtsI that generates a transient binding site for ^E^FtsN in the periplasm. This interaction is aided by ^Cyto^FtsN interacting with this FtsA in the cytoplasm ^9, 10, 31^ resulting in stabilization or clamping together of FtsL and FtsI by FtsN which generates a signal that is transmitted through FtsI to activate FtsW. Because of the dual requirement for FtsN localization to the divisome: ^E^FtsN with ^AWI^FtsL and FtsI in the periplasm and ^Cyto^FtsN with FtsA in the cytoplasm FtsN functions both as a sensor for the assembly of a functional divisome and then as its activator. Furthermore, our results reveal the roles of two key FtsL residues, L86 and E87, in the activation of FtsW.

Our revised model explains why FtsN is the last essential protein to arrive at the divisome and how multiple interactions between FtsN and the divisome contribute to its localization to the Z ring. FtsN recruitment requires ^Cyto^FtsN interaction with FtsA but also requires that FtsQLB and FtsI be present ^9, 10, 15, 32^. We propose that when FtsQLB and FtsWI are linked to FtsA and ^Cyto^FtsN interacts with this occupied FtsA it enhances the binding of ^E^FtsN to the transient interface formed between ^AWI^FtsL and FtsI resulting in a signal being transmitted to FtsW. Moreover, in light of the AF2 structure and our genetic data, we reason that in WT cells FtsI, recruited by FtsW bound to the cytoplasmic and likely the membrane domain of FtsL, contacts ^AWI^FtsL forming a weak interaction in the absence of FtsN. This transitory interaction between ^AWI^FtsL and FtsI is too weak to activate FtsW but when bound by ^E^FtsN, which stabilizes the ^AWI^FtsL-FtsI interaction, an activation signal is generated for FtsW.

Consistent with our previous model ^8^, *ftsL* and *ftsB* superfission mutations appear to enhance the ^AWI^FtsL-FtsI interaction, likely enhancing the affinity of the divisome for ^E^FtsN. In support of this, a previous study detected localization of ^E^FtsN to the divisome in the presence of *ftsB^E56A^*^16^ although it cannot be observed in a WT background ^11^. Also, consistent with the model, our previous study found that ^ΔCyto^FtsL (which is deficient in recruiting FtsWI) is readily rescued by a small increase in FtsI but not FtsW; we suggest that increasing FtsI promotes the ^AWI^FtsL-FtsI interaction to create the binding site for ^E^FtsN and rescuing recruitment ^8^. In this study we detected the localization of ^E^FtsN in a strain with the *ftsW^E289G^* superfission mutation. Thus, even though *ftsW^E289G^* bypasses FtsN our ability to detect localization of ^E^FtsN supports the idea that superfission mutations, even if *ftsW*, enhance the ^AWI^FtsL-FtsI interaction increasing the affinity for ^E^FtsN.

In this model the ^SPOR^FtsN domain is not required for the activation event and is only required for continued septal PG synthesis once the initial activation event has occurred. This is consistent with the SPOR domain binding to denuded PG that is only formed once the initial activation has occurred resulting in some septal PG synthesis and amidase action ^11, 12^. In other words, the ^SPOR^FtsN domain binding to denuded PG increases the local concentration of ^E^FtsN at the divisome thereby enhancing the ^AWI^FtsL-FtsI interaction.

In the AF2 structure ^E^FtsN is modeled to bind to ^AWI^FtsL and FtsI with the three critical residues of ^E^FtsN, W83, Y85 and L89 making hydrophobic contacts. W83 binds only to FtsL whereas Y85 and L89 bind to both FtsI and FtsL. Lineups of FtsN from various Gram-negative bacteria confirmed that hydrophobic residues are conserved at these positions with the 89^th^ position always occupied by leucine. L89 of ^E^FtsN is packed against P170 and Y168 of FtsI and Y85 of ^E^FtsN. Besides contacting L89 of ^E^FtsN, Y85 of ^E^FtsN is also packed against Y168 of FtsI. Interestingly, FtsN cannot be identified in Gram positive organisms and the ^AWI^FtsL domain is not well conserved among Gram positive bacteria suggesting some variation on the activation mechanism is likely in these bacteria ^33^.

To understand the role of FtsL in the mechanism of activation, we focused on two residues in FtsL originally identified by dominant negative mutations, *ftsL^E^*^87^ and *ftsL^L^*^86^ ^8^. Although mutation of other residues in the AWI region leads to a dominant negative phenotype, *ftsL^E87K^*and *ftsL^L86F^* are the strongest with the AF2 model indicating overlapping but distinct roles for these two residues. FtsL-L86 contacts both FtsI and FtsN and lies in a hydrophobic pocket created by 4 residues from FtsI and 2 residues from ^E^FtsN, both of which are critical ^E^FtsN residues (Fig. 2b). This pocket constrains the type of residue that can occupy this position which we confirmed by mutagenesis, as only a valine substitution, among those tested, complemented Δ*ftsL*. In contrast, FtsL-E87 is modeled to only contact the backbone of ^E^FtsN through hydrogen bonds to the primary amides of R84 and Y85 (Fig 2c). Consistent with this, previous mutagenesis revealed that only an aspartic acid residue at this position was able to restore some activity ^8^. The different interactions of these two residues of FtsL with FtsI and ^E^FtsN in the AF2 model indicate they should have separable activities.

Our study revealed *ftsL^E87K^* blocks the effect of ^E^FtsN by preventing ^E^FtsN localization. We took advantage of a strain containing the *ftsW^E289G^* superfission mutation which bypasses FtsN and suppresses *ftsL^E87K^*. Even though this allele of *ftsW* bypasses FtsN, division can be further activated as cells become shorter in the presence of FtsN or superfission mutations in *ftsL* and *ftsB*. Since *ftsW^E289G^* does not require an activation signal, it allowed construction of strains containing various alleles of *ftsL* and testing the effects of ^E^FtsN. Cells containing *ftsW^E289G^* and *ftsL* or *ftsL^E87K^*were slightly elongated (consistent with being Δ*ftsN*) whereas cells containing *ftsL* with two superfission mutations or the superfission mutations plus *ftsL^E87^*^K^ were considerably shorter. This latter result indicated that *ftsW^E289G^* was responsive to the superfission mutations and that *ftsL^E87K^* did not block the signal provided by these mutations. In addition, the strain containing *ftsL* was responsive to ^E^FtsN and became 27% shorter, whereas the strain with *ftsL^E87K^* did not. Consistent with this, *ftsL^E87K^*prevented the localization of ^E^FtsN. These results demonstrated that *ftsL^E87K^* prevented signaling from ^E^FtsN which is consistent with the AF2 model, where the only role of FtsL-E87 is to bind the backbone of the ^E^FtsN. Previously, we suggested that *ftsL^E^*^87^ prevented interaction of FtsL and FtsI ^8^. However, since *ftsL^E87K^* prevents ^E^FtsN from binding to ^AWI^FtsL-FtsI, the ^AWI^FtsL-FtsI interaction is too weak and transient (without ^E^FtsN) to be detected by the assays we used in the earlier study.

In a second approach we used superfission mutations in *ftsL* and *ftsB* to determine if the signal from ^E^FtsN goes through ^AWI^FtsL to FtsI and to FtsW. Our reasoning was that superfission mutations mimic ^E^FtsN action and that they should send a signal to FtsW unless the dominant negative mutations in *ftsL* prevent its propagation. *ftsL^G92D^* was able to suppress *ftsL^E87K^* but was unable to suppress *ftsL^L86F^* or other L86 substitutions, indicating L86 is required for signal propagation to FtsW although E87 is not. Another superfisson mutation in *ftsL*, *ftsL^E88K^*, was unable to suppress *ftsL^E87K^*but, combining it with another superfission mutation, *ftsB^E56A^*, resulted in suppression. Our previous results showed that superfission mutations are often synergistic ^8^. Nonetheless, the combination of these two superfission mutations was unable to rescue any of the substitutions at FtsL-L86 that are dominant negative. Thus, L86 (and its hydrophobic interaction with FtsI) is required to transmit the signal provided by these superfission mutations that mimic ^E^FtsN action.

Consistent with the ^AWI^FtsL-FtsI interaction being important for FtsW activation, *ftsI^Y168K^* could not complement FtsI^t*s*^ indicating it was a LOF mutation. However, it was rescued by *ftsW^E289G^*, indicating the TP activity of FtsI^Y168K^ is intact and that the interaction between ^AWI^FtsL and FtsI is not required if FtsW is already activated. In other words, the interaction is this region only plays a role in activation of FtsW. In our previous model we suggested ^E^FtsN makes ^AWI^FtsL available to interact with FtsI to activate FtsW but this can be updated to ^E^FtsN stabilizes the ^AWI^FtsL-FtsI interaction to activate FtsW. Since FtsW^E289G^ is almost fully active, fastening of ^AWI^FtsL to FtsI by FtsN is no longer necessary, even though FtsN can still enhance activation. Because FtsW^E289G^ is almost fully active it can suppress either Δ*ftsN* or *ftsL^L86F,E87K^*.

One intriguing question that remains unanswered is how ^E^FtsN’s binding to the ^AWI^FtsL-FtsI complex is transmitted to the active site of FtsW located in the periplasm proximal to the membrane. Recently, Shlosman *et al.* ^34^ proposed that an extended open conformation of PBP2 (promoted by MreC) activates RodA. Jan Lowe’s group used cryo-EM and found that *Pa*FtsI complexed with *Pa*FtsQLB exists in a partially open form with its TP domain heading away from *Pa*FtsW but short of reaching the PG layer ^35^. However, an AF2-generated model of the complex, similar to the one here, has *Pa*FtsI more extended which could reach the PG layer. Britton *et al.* used an AF2-based prediction and molecular dynamic (MD) simulations which support both our earlier model ^8^ and our revised model ^20^. In their model, like ours reported here, ^E^FtsN tethers ^AWI^FtsL to FtsI, thereby impacting the FtsW catalytic region. We can add however, that the dominant negative mutations in *ftsL* which negate the ^AWI^FtsL-FtsI interaction are suppressed by activation mutations in *ftsA* or *ftsW*. This argues the ^AWI^FtsL-FtsI interaction is not required for maintaining FtsI in the open state but is only required for activation. In the AF2 model there are multiple interactions between FtsI and FtsL (Fig. S1) and they may also promote the open state. In any event a detailed mechanism of FtsW activation must involve ^AWI^FtsL and FtsI binding (stabilized by ^E^FtsN), affecting FtsW’s catalytic region as Britton *et al.* ^20^ proposed with the anchor domain of FtsI affecting the ECL4 of FtsW..

## Materials and Methods

### Bacterial Strains and Growth Conditions

JS238 [*MC1061*, *araD* Δ(*ara leu) galU galK hsdS rpsL* Δ(*lacIOPZYA) X74 malP::lacIQ srlC::Tn10 recA1*] was primarily used for most cloning experiments, propagation and maintenance of plasmids, and dominant negative tests for *ftsL* alleles ^8^. AM1992, PK4-1 (W3110 *leu*::*Tn10 ftsL*::*kan/*pKTP108 [*repA*^ts^, *spc^R^*, P_syn135_::*ftsL*] *recA::spc*), PK23 (MCI23^ts^, *recA::spc*), SD439 (W3110 *ftsL::kan*/pDS296 [*cat*^R^, P_ara_::*ftsL*]), SD488 (W3110 *leu::*Tn*10 ftsW^E2^*^89^), BL86 [W3110 *ftsN*::*kan/*pBL200 [*repA*^ts^, spc^R^, P_syn135_::*ftsN*], Δr*ecA* Δ*lacIZYA leu*::Tn*10*], BL167 (TB28 *ftsB^E56A^*) and PK169-1 (*ftsB^E56A^*, *recA::spc, ftsL::kan*/ pDS296 [*cat*^R^, P_ara_::*ftsL*]) were previously described ^8, 30^. To construct PK498 (SD488 Δ*ftsN [ftsN::kan]*), P1 was grown on BL86 and used to transduce SD488 to Kan^R^ on LB agar plates containing 25 µg/ml kanamycin at 30°C. To eliminate the kanamycin cassette, a subcloned transductant was transformed with pCP20 [*repA*^ts^, *bla*^R^, λP_R_::*flp*] at 30°C and restreaked at 37°C for selection of clones sensitive to ampicillin and kanamycin. PK4980 (PK498 *ftsL::kan*/pSD256 [*repA*^ts^, spc^R^, P_syn135_::*ftsL*]) was generated by P1 transduction of *ftsL::kan* from PK4-1 into PK498/pSD256. Clones that grow only at 30°C, not at 37°C, and are resistant to 25 µg/ml kanamycin and spectinomycin were selectedc. PK4980-1 (W3110 *leu::Tn10 ftsW^E289G^*, Δ*ftsN*, *ftsL::kan*/pSD296-*ftsL*^WT^), PK4980-2 (W3110 *leu::Tn10 ftsW^E289G^*, Δ*ftsN*, *ftsL::kan*/pSD296-*ftsL*^E87K^), PK4980-3 (W3110 *leu::Tn10 ftsW^E289G^*, Δ*ftsN*, *ftsL::kan*/pSD296-*ftsL*^E87K,E88K,G92D^), PK4980-4 (W3110 *leu::Tn10 ftsW^E289G^*, Δ*ftsN*, *ftsL::kan*/pSD296-*ftsL*^E88K,G92D^) were created by transforming PK4980 with pSD296 (*ftsL*^WT^, pSD296-*ftsL*^E87K^, pSD296-*ftsL*^E87K,E88K,G92D^ and pSD296-*ftsL*^E88K,G92D^, respectively at 30°C and restreaking the respective colonies at 37°C. Clones were selected that were sensitive to spectinomycin and grew only in the presence of 0.2% arabinose. PK439 (SD439, *recA::spc*) was generated by P1 phage transduction of *recA::spc* from AM1992 donor to SD439 recipient at 30°C followed by selection of clones resistant to 25 µg/ml spectinomycin. PK439-2 (SD439, *recA::Tn10*) was created by P1 transduction of *srlC::Tn10 recA1* from JS238 into SD439. PK488 (SD488 *recA::spc*) was created by using the same approach as described for PK439. Unless stated otherwise, Luria-Broth (LB) medium containing 0.5% NaCl was used at indicated temperatures. For selection on LB agar and in LB broth, the following antibiotics and reagents were added at the indicated final concentrations as necessary (ampicillin,100 μg/ml; kanamycin, 25 µg/ml; chloramphenicol, 10 µg/ml; spectinomycin, 25 µg/ml; tetracycline 10 µg/ml; IPTG. 0-250 µM; glucose 0.2%; and arabinose 0.2%).

### Plasmids

All the constructs including the list of primers used in this study such as pSD256 (*repA*^ts^, spc^R^, P_syn135_::*ftsL*), pDS296 (*cat*^R^, P_ara_::*ftsL*), pMG14 *(bla^R^,* P_lac_::*^TT^gfp-ftsN*^71–^*^1^*^05^), pKTP100 (*bla^R^*, P_tac_::*ftsL*) and pKTP109 (*cat*^R^, P_ara_::*ftsI*) were previously described ^8^. All primers are available on request.

### Site-Directed Mutagenesis

Specific point mutations in *ftsL and ftsI* were introduced into pDS296 (*cat*^R^, P_ara_::*ftsL*), pKTP100 (*bla^R^*, P_tac_::*ftsL*) and pKTP109 (*cat*^R^, P_ara_::*ftsI*) by using the QuickChange site-directed mutagenesis kit according to the manufacturer’s instruction (Agilent Technologies). The list of primers used in this study was previously published.

### Alignment of the essential (E) domain of FtsN

FtsNs from a variety of Gram negative bacteria were obtained from Uniprot and aligned. Only the region around theessential (E) domain of FtsN(^E^FtsN) is depicted.

### Prediction of protein complex structures

Models of the FtsQLBWI and FtsQLBWIN complexes were produced with AlphaFold2.2 multimer ^36^ and run locally suing the full-length protein sequences. The pITM+PTM scoring metrics of the tom ranked model were 0.7019417 and 0.685166 respectively. Strctures were visualized with the PyMOL Molecular Graphics System, Version 2.5.2 Schrödinger, LLC. Structures were visualized using PyMOL (Molecular Graphics System Version 2.5.4, Schrödinger, LLC).

### Analysis of *ftsL* deletion

To confirm deletion of *ftsL* by P1 transduction, genomic DNA from W3110, PK4980-1, PK4980-2, PK4980-3 and PK4980-4 was isolated and subjected to PCR with a pair of primers used for constructing pSD256 plasmid as previously described ^8^. These primers target 250 bp upstream and downstream of the *ftsL* ORF. Amplified PCR products were analyzed by 0.8% agarose gel electrophoresis. Sequences of primers are available upon request.

### Microscopy

To assess the effects of *ftsL* mutations on cell division, cell phenotypes were monitored utilizing phase-contrast microscopy. The colonies of PK4980-1 (W3110 *leu::Tn10 ftsW^E289G^*, Δ*ftsN*, *ftsL::kan*/pSD296*-ftsL*^WT^), PK4980-2 (W3110 *leu::Tn10 ftsW^E289G^*, Δ*ftsN*, *ftsL::kan*/pSD296*-ftsL*^E87K^), PK4980-3 (W3110 *leu::Tn10 ftsW^E289G^*, Δ*ftsN*, *ftsL::kan*/pSD296*-ftsL*^E87K,E88K,G92D^), PK4980-4 (W3110 *leu::Tn10 ftsW^E289G^*, Δ*ftsN*, *ftsL::kan*/pSD296*-ftsL*^E88K,G92D^) were picked and cultured overnight at 37°C in the presence of 0.2% arabinose and chloramphenicol (34 µg/ml), kanamycin (25µg/ml) and tetracycline (10 µg/ml). Next day, overnight cultures were diluted 1/200 in the presence of the appropriate antibiotics. Samples of cells in exponential phase were immobilized on an LB agarose pad and recorded using a cooled CCD camera and processed using Metamorph (Molecular Devices) and Adobe Photoshop. To examine if FtsL^E87K^ responds to ^E^FtsN, PK4980-1 (*ftsW^E289G^*, Δ*ftsN*, pSD296-*ftsL^WT^*) and PK4980-2 (*ftsW^E289G^*, Δ*ftsN*, pSD296-*ftsL^E87K^*) were transformed with empty vector or pMG14 *(bla^R^,* P_lac_::*^TT^gfp-ftsN*^71–^*^1^*^05^) and colonies picked and cultured overnight in the presence of 0.2% arabinose, ampicillin (100 µg/ml) and chloramphenicol (34 µg/ml), kanamycin (25 µg/ml) and tetracycline (10 µg/ml) as described above. Next day, overnight cultures were diluted 1/200 in the presence of the appropriate antibiotics including 200 µM IPTG and cultured at 37°C until exponential phase. Phase-contrast images were recorded as above and cell lengths (over 200 cells per each sample) were determined with Metamorph image analysis software (Molecular Devices). For statistical analysis, one-way analysis of variance (Bonferroni method) was used with GraphPad Prism (GraphPad software).

