## Supplement file for "The essential domain of FtsN triggers cell division by promoting interaction between FtsL and FtsI"

Supplementary Figures.

Fig. S1. Alphafold model of FtsQLBWIN

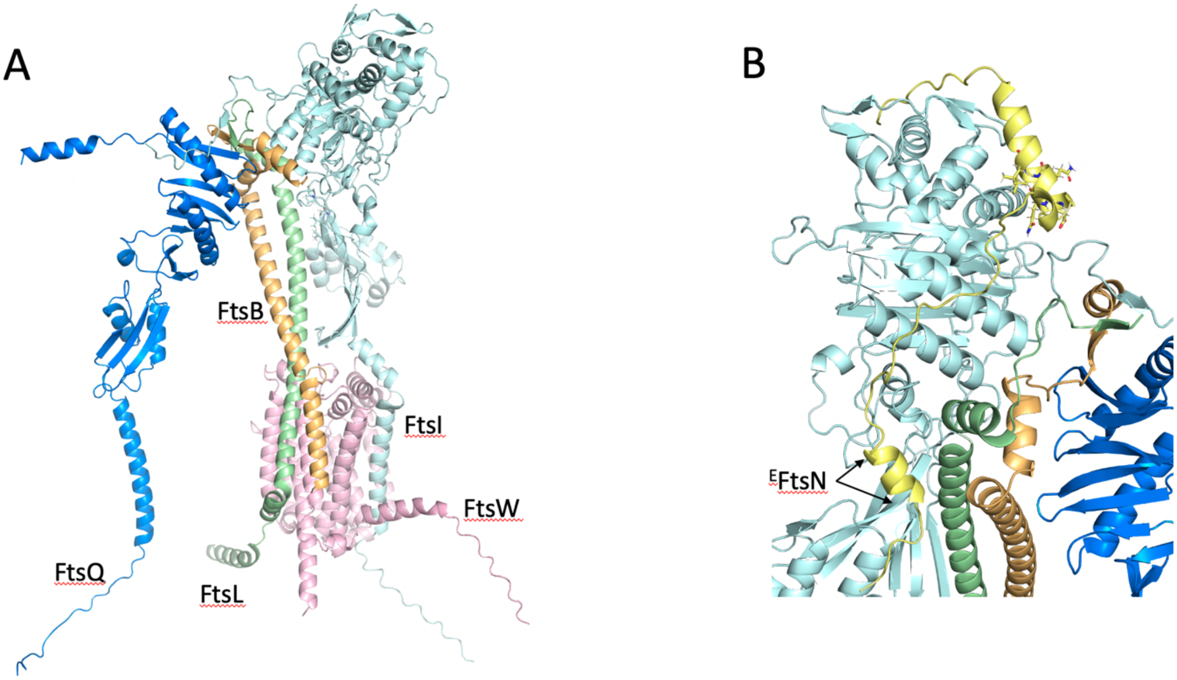


Fig. S1. AF2 multimer model of FtsQLBWI complex. A. Model of the FtsQLBWI complex was obtained using AlphaFold2.2 multimer [36]. B. Model of a 63-residue fragment (amino acids 78-141) from FtsN that binds to both FtsI and FtsL. The region corresponding to $^{E}$FtsN and residues altered in the H3 mutant are indicated.

Fig. S2. Effect of substitutions at FtsL-L86 on function

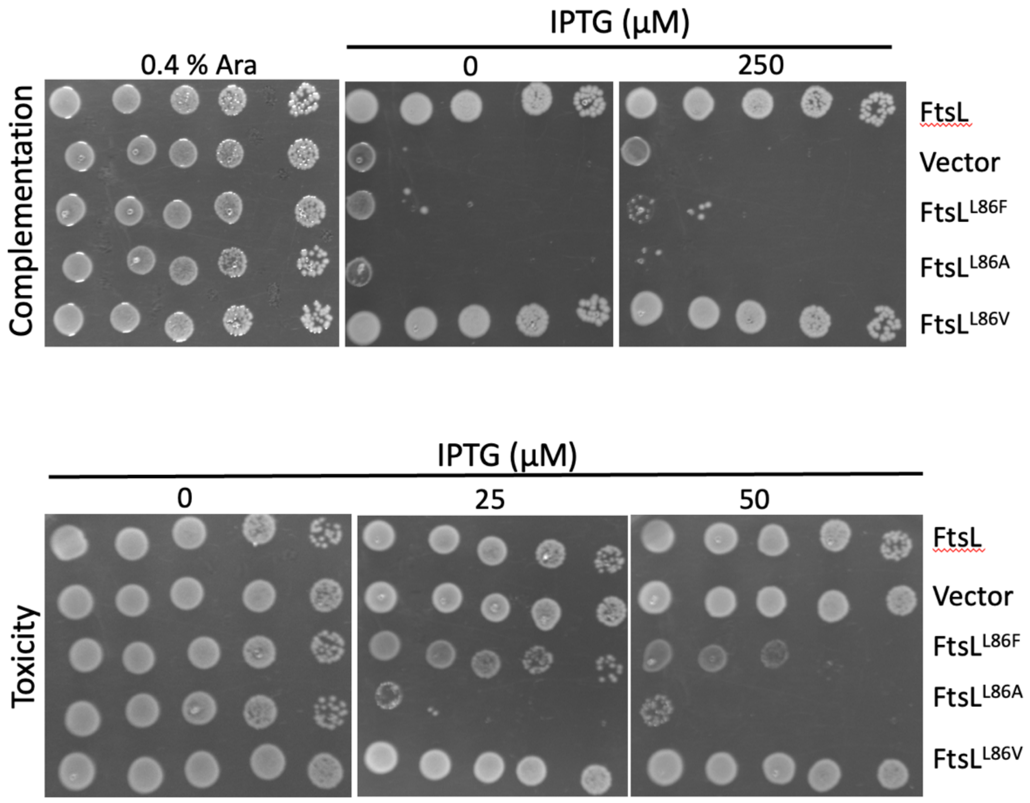


Fig. S2. Effect of various substitutions at FtsL-L86 on FtsL function. The top panel shows complementation of an FtsL depletion strain (PK439-2) by various FtsL substitution mutants. The bottom panel shows the toxicity of the same mutants as in the top panel when expressed in a WT strain (JS238).

Fig. S3. The E domain contacts FtsL in the AWI region

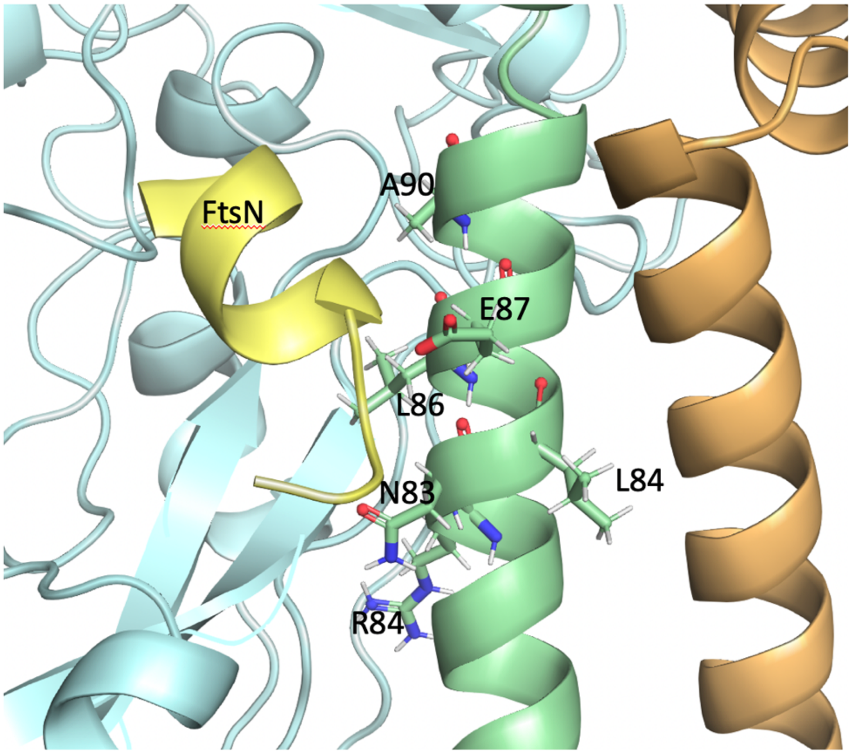


Fig. S3. Model showing the juxtaposition of $^{E}$FtsN with $^{AWI}$FtsL. The various residues that constitute $^{AWI}$FtsL (as defined by dominant negative mutations) are indicated. The 3 critical residues in $^{E}$FtsN are W83, Y85 and L89. A key FtsN residue W83 packs against hydrophobic residues in FtsL-L84 and FtsL-N83. Amides of FtsN-Y85 and R84 interact with FtsL-E87, whereas FtsN-L89 abuts FtsL-A90.

Fig. S4. Alignment of the E domain from Gram negative bacteria

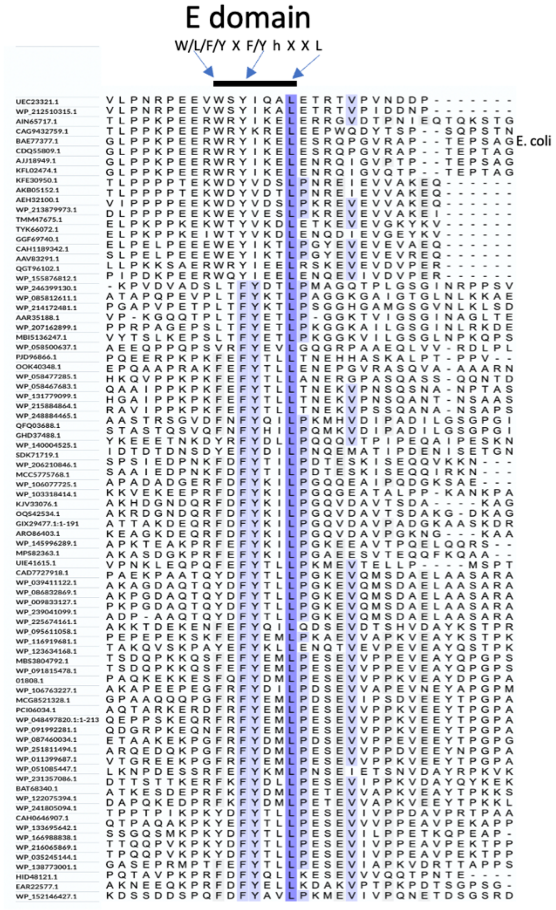


Fig. S4. Alignment of $^{E}$FtsN from Gram-negative bacteria. FtsN sequences from a wide range of Gram-negative bacteria were aligned and the region around $^{E}$FtsN is shown with the consensus sequences displayed at the top (W/L/F/Y X F/Y h X X L; where h stands for large hydrophobic residue). Although many bacteria have FtsN, we could not identify one in *Neisseria*. Interestingly, *Caulobacter* has a gene designated *ftsN* but we could not identify an E domain.

Fig. S5. Cluster of charged residues

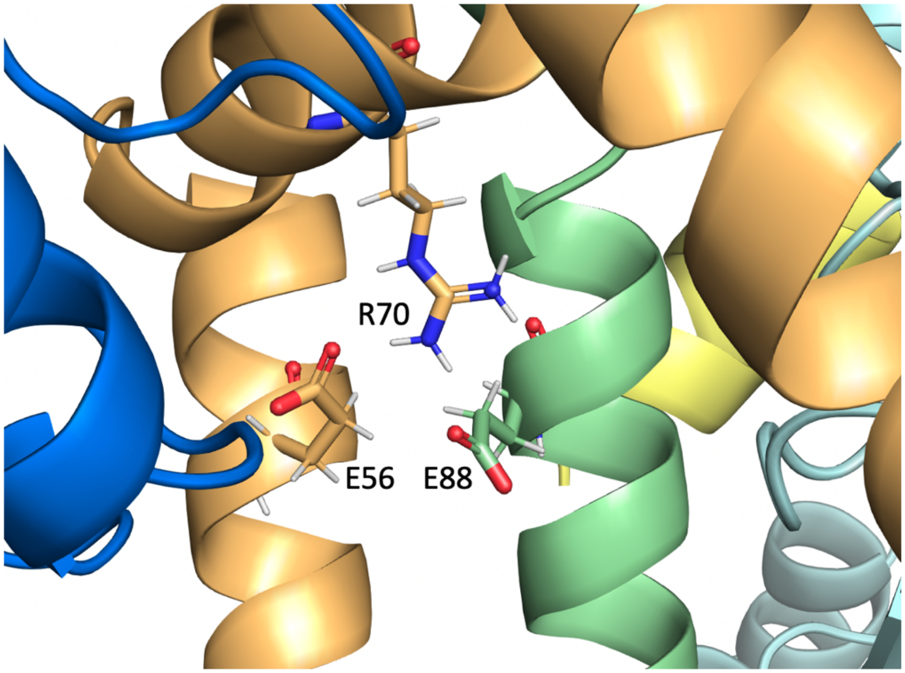


Fig. S5. The CCD domains of FtsL and FtsB contain a cluster of charged residues. A cluster of three charged residues are present in the CCD domains of FtsL and FtsB. They include B56 and R70 from FtsB and E88 from FtsL. Eliminating the charge of these residues produces superfission mutations.

Fig. S6. A second region of FtsN interacts with FtsI

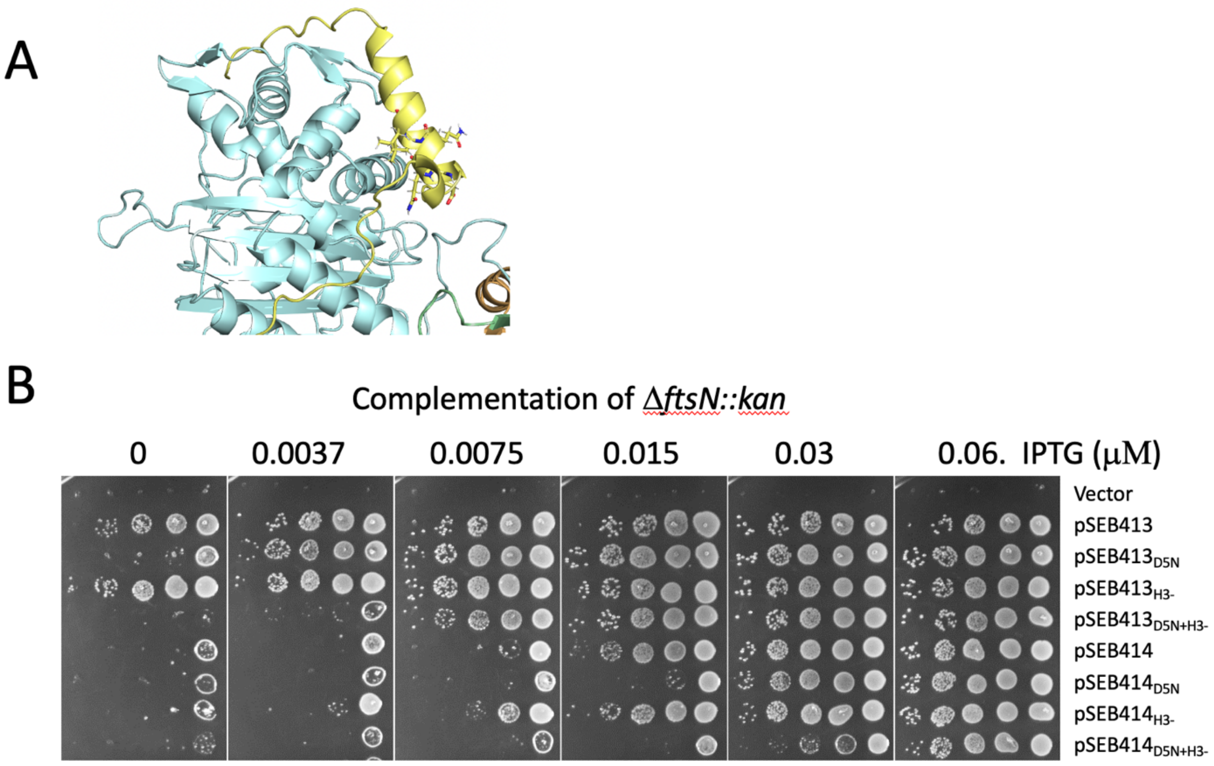


Fig. S6. A second region of $^{E}$FtsN interacts with FtsI. A) A region of FtsN which overlaps with putative helix 3 contacts FtsI. Three conserved residues (E117A Q118, Q120) were mutated to alanine, designated H3 and tested for complementation. B) Complementation test of the triple H3 mutant. The triple mutant (H3) was tested in full length FtsN and FtsN$^{1\text{-}140}$. In addition, these constructs contained either WT FtsN or FtsN$^{D5N}$. The FtsN$^{D5N}$ mutant requires more IPTG to complement than WT and the addition of H3 to the Dr mutant required even more IPTG to complement. The plasmids are pSEB413 (*ftsN*), and pSEB414 (ftsN$^{1\text{-}140}$). These plasmids contained the H3 mutations, D5N or both. The parent vector is pDSW210.

Fig. S7. EFtsN binding to FtsQLBWI brings FtsL and FtsI closer together

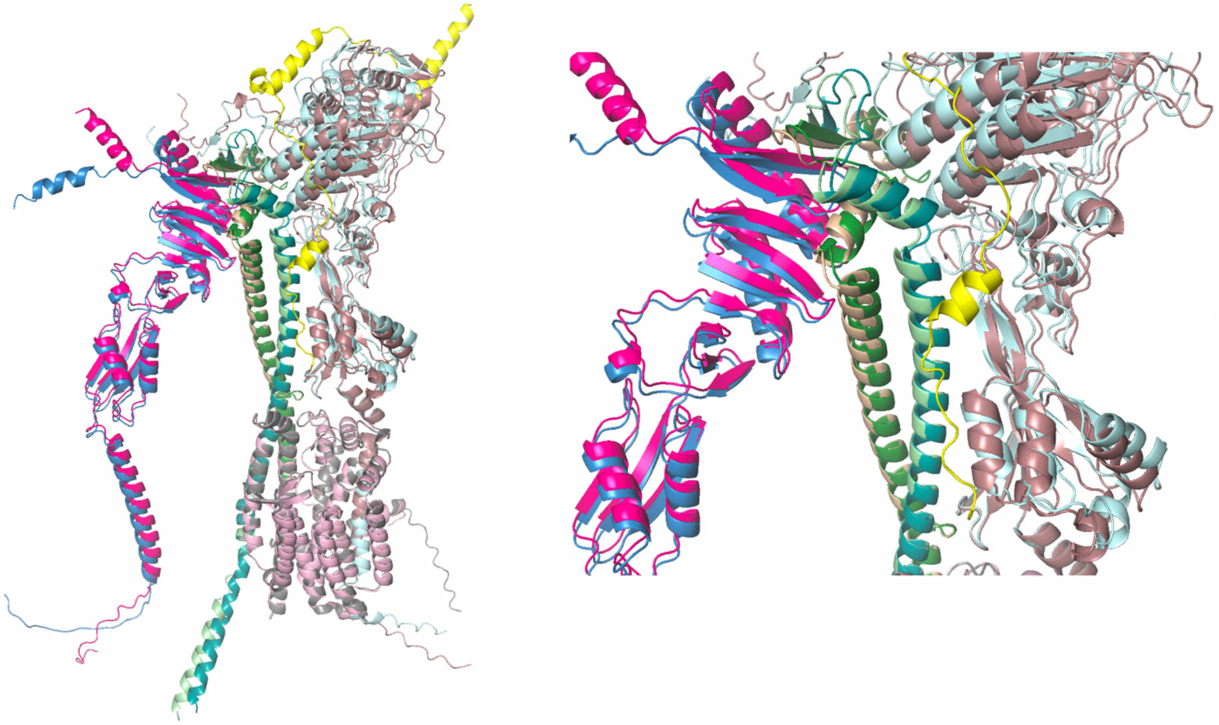

Fig. S7. EFtsN binding to FtsQLBWI brings FtsL and FtsI closer together. The FtsQLBWIN model was superimposed on the FtsQLBWI model. The FtsQLBWI model is colored and in Fig. S1A. In the model with FtsN the proteins have a darker tint. I For FtsL P112 to FtsI I486 the distance goes from 8.1 down to 6.8 Å when FtsN is added.

### Fig. S8. Combining *ftsL*$^{G92D}$ and *ftsL*$^{E88K}$ weakly bypasses FtsN

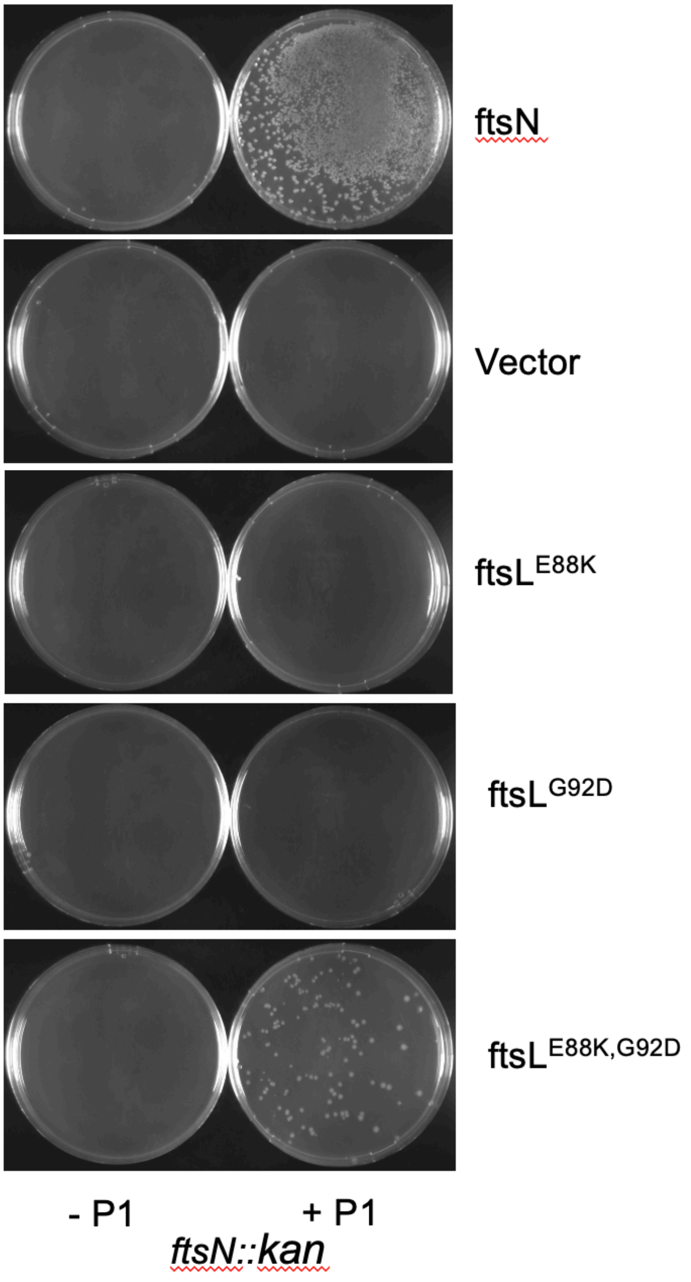


Fig. S8. Combining *ftsL*$^{G92D}$ and *ftsL*$^{E88K}$ in full length *ftsL* weakly bypasses *ftsN*. W3110 *leu*::Tn*10* derivatives containing various plasmids were infected or not with P1 phage grown on a strain containing *ftsN::kan*. The plasmids contained *ftsN* or alleles of *ftsL*

under *lac* promoter control. Transductants were selected on plates containing antibiotics and 100 μM IPTG. Transductants were obtained if the plasmid contained *ftsN* or *ftsL*$^{E88K,G92D}$ but not if the plasmid contained *ftsL* alleles with a single mutation. Transductants were obtained in the presence of *ftsL*$^{E88K,G92D}$ indicating *ftsN* was bypassed but cells displayed elongated morphology indicating that *ftsN* was weakly bypassed.

Fig. S9. Characterization of strains containing different FtsL alleles.

A

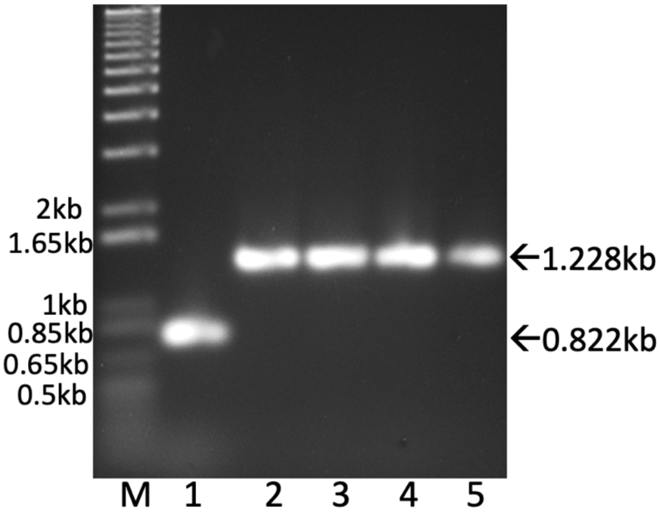


B

M-1kb plus DNA ladder
1-W3110 Wild Type
2-PK4980-1 (W3110 *leu::tn10*, *ftsW*$^{E289G}$, *ftsN* < > *frt*, *ftsL*::*kan*/pSD296[pBAD33 *ftsL*])
3-PK4980-2 (W3110 *leu::tn10*, *ftW*$^{E289G}$, *ftsN* < > *frt*, *ftsL*::*kan*/pSD296[pBAD33 *ftsL*$^{E87K}$])
4-PK4980-2 (W3110 *leu::tn10*, *ftsW*$^{E289G}$, *ftsN* < > *frt*, *ftsL*::*kan*/pSD296[pBAD33 *ftsL*$^{E87K,\ E88K,\ G92D}$])
5-PK4980-2 (W3110 *leu::tn10*, *ftsW*$^{E289G}$, *ftsN* < > *frt*, *ftsL*::*kan*/pSD296[pBAD33 *ftsL* $^{E88K,\ G92D}$])

Fig. S9. Characterization of strains of Δ*ftsN ftsW*$^{E289G}$ containing different *ftsL* alleles. A) PCR analysis of WT and Δ*ftsL* strains (derivatives of PK4980 (W3110 *leu*::Tn*10 ftsW*$^{E289G}$, Δ*ftsN, ftsL::kan*]) containing plasmids with different *ftsL* alleles under arabinose promoter control (from Fig. 3). The PCR fragment from the strains with *ftsN::kan* is larger due to the presence of the *kan* in place of *ftsN*. The plasmids are derivatives of pSD296 ($P_{BAD}$::*ftsL*). B) Phenotype of Δ*ftsN ftsW*$^{E289G}$ strains carrying various *ftsL* alleles.
